# DNA barcoding and olfactory identification of attractive nectar sources for *Aedes aegypti* mosquitoes

**DOI:** 10.64898/2026.06.19.733381

**Authors:** Sandeep Jandu, Anand Patil, Jae Paik, Mba-Tihssommah Mosore, Daniel Kline, Edmund J. Norris, Edwin R. Burgess, Jeffrey A. Riffell

## Abstract

Adult mosquitoes rely on plant-derived sugars for survival, reproduction, and flight, yet the plant taxa that mosquitoes encounter in nature and the odors that make those plants attractive remain poorly understood. Most studies of mosquito attraction to plant odors have focused on candidate plants selected a priori, rather than plants linked to field-collected mosquitoes. Here, we combined plant DNA barcoding, semi-field behavioral assays, and volatile profiling to identify field-associated plant resources relevant to *Aedes aegypti*. Plant DNA recovered from mosquitoes collected across three Florida counties revealed broad plant associations, including 90 genera spanning 37 families, with several taxa recurring across counties or appearing prominently within particular localities. Behavioral experiments in semi-field sticky-trap assays found that five field-associated plant taxa were significantly attractive relative to blank controls, indicating that taxa associated with mosquitoes in nature can also function as attractive cues under semi-field conditions. GC-MS analyses of headspace collections from 42 plant taxa detected 211 volatile compounds and revealed substantial variation in both total emission rate and odor composition among taxa. Although several compounds, including α-pinene, limonene, 4-ethylacetophenone, 2-ethyl-1-hexanol, 4-ethylbenzaldehyde, and caryophyllene, were broadly distributed across plant groups, volatile profiles differed significantly among taxa and shared compounds often occurred at markedly different proportional abundances. The five behaviorally tested taxa likewise showed both overlap and divergence, sharing 17 compounds across all five taxa while differing in dominant constituents and total emissions. Together, these results show that *Ae. aegypti* interacts with a diverse set of plants in the field, and suggests nectar-seeking is shaped not simply by plant identity or total odor abundance, but by the composition and proportional structure of plant odors.

## Introduction

Mosquitoes rely on plant-derived sugars as an important source of energy throughout their adult lives, with both males and females regularly feeding from plants (Foster, 1995; Sissoko et al., 2019; Fikrig et al., 2020). Plant sugar feeding influences multiple aspects of mosquito biology, including survival, flight activity, mating, fecundity, and host-seeking behavior (Foster, 1995; Manda et al., 2007; Stone et al., 2012; Njoroge et al., 2021). In addition to supporting basic metabolic demands, plant-derived sugars can influence dispersal capacity and reproductive success, making access to sugar sources an important component of mosquito ecology. To locate these resources, mosquitoes use olfactory cues emitted by plants, including volatile organic compounds (VOCs) released from floral and vegetative tissues. Studies have shown that mosquitoes are attracted to plant-associated odors and that plant-derived volatile odors can influence mosquito behavior (Nyasembe and Torto, 2014; Nyasembe et al., 2012; Jhumur et al., 2008; Lahondère et al., 2020; Pullmann-Lindsley et al., 2024). However, compared with the extensive body of work examining host-seeking and blood-feeding behavior, mosquito interactions with plants and the sensory cues underlying plant-seeking behavior remain comparatively understudied.

Plant-associated odors are highly diverse, varying substantially among plant species in both composition and abundance of volatile compounds (Raguso, 2008). Floral and vegetative tissues can emit complex odors consisting of dozens to hundreds of VOCs, generating chemically distinct odor environments encountered by mosquitoes while foraging (Jhumur et al., 2008; Nyasembe and Torto, 2014; Sobhy and Berry, 2024). Correspondingly, mosquitoes do not respond equally to all plants, and studies have shown that certain flowers, fruits, and vegetative odors can elicit stronger attraction than others (Jhumur et al., 2008; Müller et al., 2011; Nyasembe et al., 2012; Kashiwagi et al., 2022). In *Aedes aegypti*, olfactory responses to plant-associated compounds have been implicated in the location of sugar sources and other plant-associated resources (Fikrig et al., 2017; Jové et al., 2020; Pullmann-Lindsley et al., 2024). At the same time, the ecological relevance of many experimentally tested plant odors remains unclear, as behavioral and chemical studies often focus on candidate plant species without evidence that mosquitoes naturally interact with those plants under field conditions. Many attraction assays are also performed under laboratory conditions that may not fully reflect the complexity of odor environments encountered outdoors. As a result, relatively little is known about which plant-associated odor sources are most relevant to mosquito plant-seeking behavior in natural environments.

Plant DNA recovered from field-collected mosquitoes provides an opportunity to characterize plant-derived material associated with naturally occurring mosquito–plant interactions and to identify plant taxa that may represent ecologically relevant sources of plant-derived sugars. Previous studies have used plant DNA recovered from mosquitoes to infer plant associations and feeding behavior, revealing that mosquitoes can interact with a broad diversity of plant taxa in natural environments (Nyasembe et al., 2018; Kinya et al., 2024; Mosore et al., 2026). These approaches provide a means of linking mosquito behavior to naturally encountered plant resources rather than relying exclusively on candidate plant species selected *a priori*. However, comparatively few studies have linked field-derived plant detections with downstream behavioral and chemical analyses to evaluate whether detected plant taxa are associated with mosquito attraction or distinct volatile profiles (Nyasembe et al., 2018; Kinya et al., 2024; Cooper et al., 2025). In addition, the extent to which plant associations vary geographically, and whether recurring plant taxa detected from mosquitoes correspond to behaviorally relevant odor sources, remains incompletely understood. Integrating molecular, behavioral, and chemical approaches may therefore provide a more ecologically grounded framework for identifying plant-associated odor sources relevant to mosquito plant-seeking behavior.

In this study, we used plant DNA barcoding to characterize plant taxa associated with field-collected *Aedes aegypti* mosquitoes from multiple counties in Florida. We then selected representative plant taxa detected in mosquito samples for semi-field behavioral assays to evaluate mosquito attraction under controlled outdoor conditions. In parallel, we characterized volatile profiles from plant taxa included in the study using dynamic headspace collections and gas chromatography–mass spectrometry (GC-MS). Together, this integrative approach provides insight into ecologically relevant plant-associated odor sources potentially involved in mosquito plant-seeking behavior.

## Materials and Methods

### Mosquito collection and sample pooling

Field-collected *Aedes aegypti* mosquitoes were obtained through collaborations with mosquito control districts in Florida. Mosquitoes were collected at multiple sites within St. Johns County, Volusia County, and Miami-Dade County during June–August 2024 using standard mosquito surveillance and control trapping protocols employed by each district. Collected specimens were predominantly female. Mosquitoes were processed in pools of 1–3 individuals, with most samples consisting of three mosquitoes per pool. Pools were composed of mosquitoes of the same sex collected from the same trap on the same collection day.

### Surface sterilization and DNA extraction

Prior to DNA extraction, mosquitoes were surface-sterilized by rinsing vigorously with 70% ethanol under light vacuum. Mosquitoes were transferred into bead-beating tubes using ethanol-sterilized forceps. DNA was extracted from whole mosquitoes using a hybrid CTAB–silica column protocol. Mosquitoes were homogenized using a Mini-Beadbeater-16 (BioSpec Products) with 1-mm zirconium beads for 1 min in CTAB lysis buffer (100 mM Tris-HCl, 20 mM EDTA, 1.4 M NaCl, 2% (w/v) CTAB, 0.2% (v/v) 2-mercaptoethanol). Homogenates were incubated overnight at 65 °C, followed by Proteinase K treatment at 56 °C. DNA was separated using two sequential extractions with phenol:chloroform:isoamyl alcohol (25:24:1). The aqueous phase was purified using the Qiagen DNeasy Blood & Tissue Kit according to the manufacturer’s instructions beginning at the binding step. DNA was eluted in 200 µL Buffer AE and stored at −20 °C.

### Nested PCR amplification of plant DNA

Plant DNA was amplified using a two-step nested PCR approach targeting the chloroplast *rbcL* gene to increase sensitivity and specificity of amplification from low-concentration plant DNA. The first round of PCR was performed using outer primers (5′-CACCACAAACAGAGACTAAAGC-3′ and 5′-TCTCTCCAACGCATAAACGG-3′) with an annealing temperature of 54 °C. Nested amplification was subsequently carried out using 1 µL of first-round PCR product as template. The second round employed inner primers (5′-ACTAAAGCAAGTGTTGGATTCAAAG-3′ and 5′-GTAAAATCAAGTCCACCRCG-3′) with an annealing temperature of 58 °C. PCR products were visualized on agarose gels to confirm successful amplification prior to downstream sequencing.

### Nanopore library preparation and sequencing

Nanopore sequencing was performed using Oxford Nanopore Technologies native barcoding protocols with R10.4.1 chemistry and the Native Barcoding Kit 96 (V14; Oxford Nanopore Technologies, Oxford, UK). PCR products from successful nested amplifications were transferred to a 96-well plate format for library preparation. For each sample, PCR product was combined with nuclease-free water, and negative control wells containing only nuclease-free water were included on each plate to monitor for potential barcode misassignment during sequencing. Amplicons were subjected to end repair and dA-tailing using the NEBNext Ultra II End Repair/dA-Tailing Module (New England BioLabs, Ipswich, MA, USA) according to the manufacturer’s instructions. Individual samples were then ligated to unique barcode sequences using the Native Barcoding Kit and NEB Blunt/TA Ligase Master Mix (New England BioLabs). Barcoded samples were pooled into a single library and cleaned according to the Oxford Nanopore Technologies protocol. Sequencing adapters were ligated to the pooled library using the NEBNext Quick Ligation Module (New England BioLabs). The adapter-ligated library was washed, eluted in elution buffer, and quantified prior to sequencing. Libraries were normalized to an appropriate loading concentration and combined with sequencing buffer and loading beads according to the manufacturer’s instructions before loading onto R10.4.1 flow cells.

Sequencing was performed on a MinION device using MinKNOW software, and raw electrical signal data were basecalled using Dorado. Initial quality control was performed during basecalling using a minimum quality score threshold of Q ≥ 10. Reads meeting this threshold were assigned to barcode-specific bins and exported as FASTQ files. To reduce barcode misassignment, demultiplexed reads were subjected to additional filtering requiring a barcode match of >97% and a minimum barcode alignment length of 37 bases at each end of the read, consistent with recent Nanopore metabarcoding workflows for mosquito-plant samples (Mosore et al., 2026).

### Sequence processing and taxonomic assignment

Basecalled and demultiplexed reads in FASTQ format were processed for taxonomic identification of plant DNA. Reads meeting initial quality control thresholds (minimum Q score ≥ 10) were retained for downstream analysis. Filtered reads were aligned to a reference chloroplast rbcL gene sequence from Pinus ponderosa (NCBI Accession: KC156882) using Minimap2, following the reference-based rbcL filtering strategy used by Mosore et al. (2026). Reads successfully mapping to the reference were retained and converted to FASTA format. Sequences were queried against the NCBI nucleotide database using BLASTn to assign plant taxonomic identities. BLAST results were further processed using a custom analysis pipeline implemented in Python to apply additional stringency to taxonomic assignments. Matches were subjected to secondary quality filtering, retaining only alignments with a minimum percent identity of 98%, alignment length ≥550 bp, bitscore ≥400, and E-value ≤1 × 10⁻²⁰.

For each sample, filtered hits were aggregated by taxon. To reduce the influence of low-frequency or spurious assignments, taxa within each sample were ranked by the number of supporting reads, and a sample-specific threshold was applied based on the 75th percentile of read counts. Taxa were retained if they met or exceeded this threshold or represented the most abundant taxon within the sample, with a minimum retention threshold of 10 reads. Filtered taxonomic assignments were then linked to sample metadata for downstream analyses.

### Plant volatile headspace collections and GCMS analyses

To characterize plant volatile profiles for subsequent chemical analysis, dynamic headspace collections were performed in Gainesville, Florida during July–September 2024 and July–September 2025, a commonly used approach for sampling floral and mosquito-associated plant volatiles (Jhumur et al., 2008; Nyasembe et al., 2018; Raguso, 2008). Plant taxa used for volatile profiling were selected to represent an ecologically informed set of candidate mosquito-associated odor sources. Selection emphasized taxa detected in preliminary mosquito-associated plant DNA surveys where possible, taxa previously reported as mosquito-attractive or otherwise relevant to mosquito plant use, and locally common plants with floral or vegetative traits consistent with potential mosquito sugar resources (Peach and Gries, 2019, 2020; Shannon et al., 2024; Upshur et al., 2023). Plant material was collected from multiple field sites across the city and transported indoors, where all volatile sampling was conducted under controlled conditions. Depending on the species, headspace was collected from whole intact plants, cut flowering stems, or vegetative tissue obtained from either field-grown plants or plants purchased from local nurseries. All collections were performed on flowering plants using fresh, intact tissue, and plant species were identified in the field at the time of collection based on morphological characteristics.

For each collection, plant material was enclosed in a PET oven bag (WRAPOK, China) secured around the stem or base of the tissue. Two polytetrafluoroethylene (PTFE) tubes were inserted at the base of the bag: one supplying charcoal-filtered air into the enclosure and the other pulling air out through an adsorbent trap. Airflow was maintained at a constant rate of 1 L min⁻¹ using a diaphragm pump (Gast 10D1125-101-1052, Gast Manufacturing, Benton Harbor, MI, USA). Headspace volatiles were collected on adsorbent traps constructed from borosilicate Pasteur pipettes (VWR, Radnor, PA, USA) containing 100 mg of Porapak Q (80/100 mesh; Waters Corporation, Milford, MA, USA). Collections were conducted at consistent start times and run continuously for 12 h per sample. The number of biological replicates varied by species, with at least three replicates collected per species.

Prior to use, Porapak Q traps were cleaned with sequential methanol and hexane washes. Immediately following collection, traps were eluted with 600 µL of hexane (≥99% purity; Sigma-Aldrich, St. Louis, MO, USA), and eluates were transferred to glass autosampler vials. Samples were initially stored at −20 °C and subsequently transferred to −80 °C for long-term storage until analysis. Field blanks were collected concurrently as headspace controls by running Porapak Q traps in empty oven bags under identical airflow and collection conditions; these controls were eluted, stored, and processed in parallel with plant samples.

For chemical analysis, 3 µL of each sample was injected into an Agilent 7890A gas chromatograph coupled to a 5975C mass selective detector (Agilent Technologies, Santa Clara, CA, USA). Volatile compounds were separated using a DB-5MS column (J&W Scientific, Folsom, CA, USA; 30 m × 0.25 mm × 0.25 µm) with helium as the carrier gas at a constant flow rate of 1 mL min⁻¹. The GC oven temperature program was as follows: 45 °C for 4 min, followed by a ramp of 5 °C min⁻¹ to 150 °C, then 15 °C min⁻¹ to 300 °C, with a final hold of 2.5 min, for a total run time of 37.5 min.

Chromatographic peaks were manually integrated using ChemStation software (Agilent Technologies, Santa Clara, CA, USA) and tentatively identified by comparison to mass spectra in the NIST library. Compound identities were further evaluated using Kovats retention indices (KRI), and data were processed to remove potential contaminants and consolidate chemical synonyms prior to downstream analyses. Emission rates were estimated from peak areas and standardized to the 12-h collection period.

### Semi-field behavioral assays and anthrone assays

Semi-field behavioral assays were conducted from July 28 to August 8, 2025 at the USDA-ARS facility in Gainesville, Florida, within outdoor screen house enclosures (height: 104 in; side length: 116 in) under ambient environmental conditions (temperature: 24–37 °C; relative humidity: 42–91%; variable cloud cover; minimal rainfall). Weather-adjusted negative-binomial GLMMs indicated that ambient conditions contributed to day-to-day variation in trap capture (χ²(3) = 18.55, p < 0.001), although plant treatment effects remained significant after weather adjustment (χ²(5) = 50.90, p < 0.001). Two independent enclosures were used per day, with each enclosure representing a single trial. *Aedes aegypti* (Orlando strain) mosquitoes were reared under laboratory conditions and used for all experiments. Cohorts consisted of a mix of males and females at approximately equal numbers and were sugar-starved for 12–24 h prior to release. A total of 300 mosquitoes were released into each enclosure at the start of each trial.

Mosquitoes were released into each enclosure at 09:30, and traps were collected at 18:00. Assays were conducted across 12 days, with two enclosures per day, for a total of 24 independent trials. The assays focused on five taxa: *Bidens alba*, *Lantana camara*, *Richardia scabra*, *Solanum lycopersicum*, and *Sorghum* sp. These taxa were selected because they were identified in preliminary surveys of mosquito-associated plant DNA, were common or readily available at field sites, and together spanned five plant families. Four of the five taxa were represented in the final filtered DNA barcoding dataset, whereas Sorghum sp. was included based on preliminary DNA barcoding results.

Behavioral assays were conducted using custom sticky traps constructed from black hardware net (Tenax 084075, Tenax Corporation, Baltimore, MD, USA) coated with adhesive (Tangle-Trap, Tanglefoot Company, Grand Rapids, MI, USA). Traps were formed into cylindrical structures (approximately 12 inches in diameter and 24 inches in height) that surrounded plant material while remaining open at the top. Six traps were placed in each enclosure, positioned 18 inches apart on the ground. Five traps were baited with cut plant material (including inflorescences, stems, and leaves) placed in glass jars containing water and cut flower food, with the opening covered in aluminum foil to prevent mosquito access to the liquid. A bamboo rod placed in an identical jar served as the control. The same plant species were used in both enclosures each day and arranged in a different randomized order. Trap locations were rotated by one position between trials to account for spatial effects. After collection, mosquitoes adhered to traps were removed manually using forceps. Removal from the adhesive frequently resulted in physical damage to specimens, which precluded reliable determination of sex; consequently, sex was not recorded. Following each behavioral assay, BG traps (BG-Sentinel traps, Biogents AG, Regensburg, Germany) were placed in each enclosure in standardized positions and operated for 1 h to collect mosquitoes that had not adhered to sticky traps. Captured mosquitoes were immediately frozen and stored until completion of all behavioral trials.

Sugar feeding was assessed using a warm anthrone assay performed on whole mosquitoes. Because mosquitoes adhered to sticky traps were unsuitable for chemical analysis, only mosquitoes collected from BG traps were used for this assay. The proportion of sugar-fed individuals among these mosquitoes was used as an indicator of recent sugar feeding activity within each trial. Mosquitoes collected from BG traps on Day 3, Day 8, and Day 12 were processed individually after completion of the behavioral trial, with processing limited to a subset of trial days, and sampling timepoints selected to span the duration of the experiment. Individual mosquitoes were homogenized in sodium sulfate buffer and processed according to standard anthrone protocols, followed by heating at 95 °C to develop the colorimetric reaction and measurement of absorbance at 625 nm using a spectrophotometer. A mosquito was classified as sugar-fed if its absorbance reading exceeded two standard deviations above that of starved negative controls. Negative controls consisted of sugar-starved mosquitoes from the same cohort that were not released and were frozen directly. A glucose standard curve was used for quantification.

### Statistical analyses

Statistical analyses were performed in Python 3 and R 4.4.3. Differences in plant taxonomic composition among counties were evaluated using permutational multivariate analysis of variance (PERMANOVA) based on Jaccard distance matrices calculated from presence/absence data at both the genus and family-only levels.

Variation in volatile composition among plant groups was evaluated using PERMANOVA and analysis of similarity (ANOSIM) based on Bray–Curtis dissimilarities calculated from relative compound proportions. Principal component analysis (PCA) was used to visualize patterns in volatile composition and identify compounds contributing to variation among samples.

For semi-field behavioral assays, sticky-trap mosquito counts were analyzed using negative binomial regression, with plant species included as the fixed effect of interest and experimental day included as a fixed blocking term. Overall treatment effects were evaluated using likelihood-ratio chi-square tests. Individual plant treatments were then compared against the blank control, with effect sizes reported as incidence rate ratios (IRRs) and associated 95% confidence intervals. P-values for plant-versus-control comparisons were adjusted using Bonferroni correction. As a secondary analysis, ambient weather variables were incorporated using negative-binomial generalized linear mixed models (GLMMs). In these models, plant species, weather covariates, and trap position were treated as fixed effects, while day and trial were treated as random effects. For anthrone assays, mosquitoes were classified as sugar-fed or non-sugar-fed based on absorbance values relative to sugar-starved controls, and sugar-feeding rates were summarized descriptively across sampled days.

## Results

### Plant taxa detected in mosquito samples

Plant-derived DNA was detected in mosquito samples from Miami-Dade, St. Johns, and Volusia counties, representing a broad range of angiosperm taxa. Across 175 samples, detections spanned 38 unique plant families. All reported taxa were supported by county-level plant availability data compiled from Florida Plant Atlas and GBIF datasets, an approach consistent with recent efforts to cross-reference mosquito-associated plant detections against Florida plant-voucher records (Mosore et al., 2026); genus-level detections were restricted to taxa present in these datasets, while detections lacking genus-level support were reported at the family level when the corresponding family was documented. Accordingly, 90 genera spanning 37 families were identified at the genus level, along with an additional 22 taxa reported at the family level only. Several plant families were consistently detected across counties, including Asteraceae, Solanaceae, Lauraceae, and Cannabaceae, while others were detected in only a subset of samples or were restricted to individual counties. The number of detected taxa varied among counties, with Miami-Dade samples containing 62 genera and 17 family-level detections, Volusia samples containing 66 genera and 15 family-level detections, and St. Johns samples including fewer taxa overall, with 20 genera and 9 family-level detections.

### Prevalence and distribution of detected taxa

Plant taxa differed in how frequently they were detected across samples, with some taxa appearing in a substantial proportion of samples and others detected only occasionally (Table 1). At the genus level, several taxa were present in a relatively large fraction of samples within individual counties. In Volusia, *Datura* was detected in 72% of samples. In St. Johns, both *Datura* and *Smilax* were among the more prevalent genera, detected in 75% and 42% of samples, respectively. In Miami-Dade, detections were more evenly distributed across genera, with no single genus occurring at similarly high frequency compared with the most prevalent genera in Volusia and St. Johns, a pattern also reflected in the more uniform distribution of prevalence values across genera (Table 1). At the family level, prevalence patterns also differed within and among counties, with Chenopodiaceae present in 77% of samples in Miami-Dade, Solanaceae detected in 74% of samples in Volusia, and Solanaceae detected in all samples (100%) in St. Johns. Several taxa selected for semi-field behavioral assays, including *Bidens*, *Lantana*, *Solanum*, and *Richardia*, were among those detected in the barcoding dataset (Table 1) and were chosen to represent taxa observed across samples. *Sorghum* sp. was included in the behavioral assays because it was detected in preliminary analyses of the barcoding dataset, although it was not retained in the final filtered dataset. Although the identity of the most prevalent taxa differed among counties, each was characterized by a small number of relatively common taxa alongside a broader set of less frequently detected taxa.

**Table 1.**
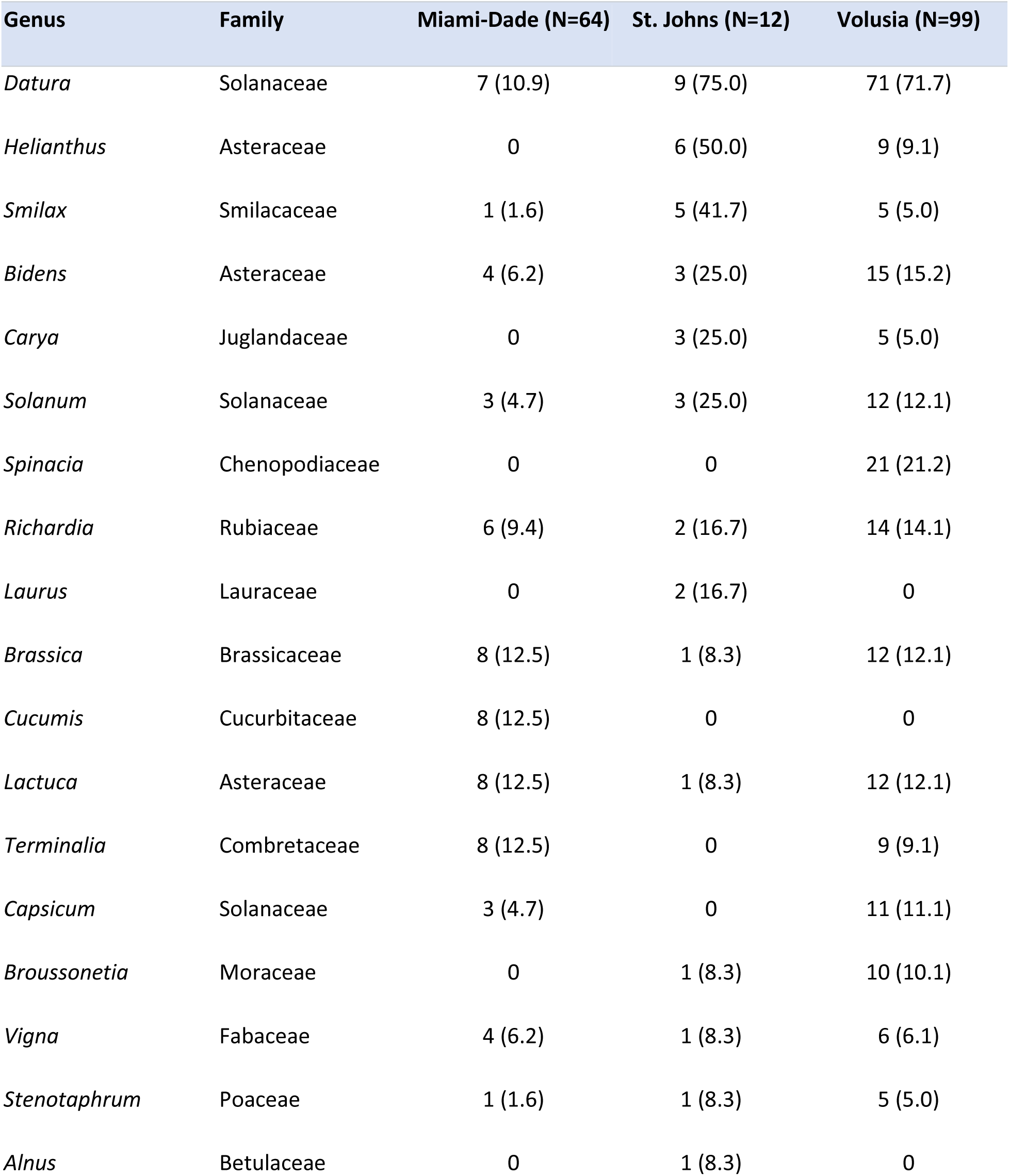
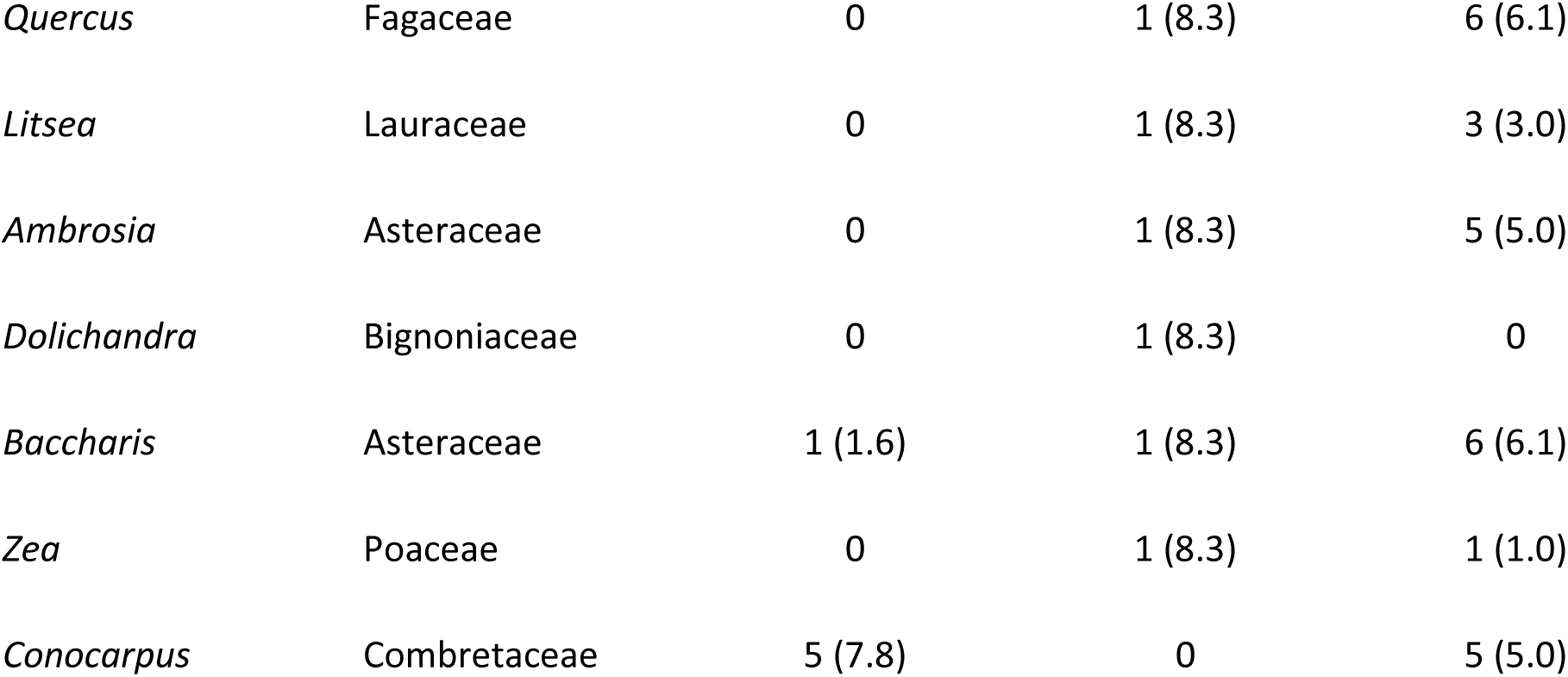
Prevalence of the top 25 plant genera detected in mosquito samples across counties. Values represent the number of mosquito samples in which each plant genus was detected, with the corresponding prevalence percentage shown in parentheses, for Miami-Dade, St. Johns, and Volusia counties based on presence/absence data.

### Differences in taxonomic composition among counties

Taxonomic composition varied among counties, with Miami-Dade and Volusia samples including a broader range of detected taxa, and most taxa occurring in a subset of samples rather than across all samples. In contrast, St. Johns County samples were characterized by a more limited set of taxa and more frequent detection of the same taxa across samples. These differences in composition were supported by PERMANOVA analyses based on Jaccard distances, which indicated significant differences among counties at both the genus level (pseudo-F = 5.13, R² = 0.059, p = 0.001) and the family-only level (pseudo-F = 29.16, R² = 0.283, p = 0.001), with a stronger effect observed at the family level. Despite these differences, there was clear overlap among counties, with several plant families detected across multiple locations, including Solanaceae, Asteraceae, Rubiaceae, and Lauraceae, indicating shared components of the detected plant community, while differences in both the number of detected taxa and their distribution across samples contributed to distinct profiles for each county, as reflected in the distribution of high-prevalence genera across counties (Table 1). Some of this variation likely reflects differences in sampling effort, but overall patterns in taxonomic composition and prevalence were consistent within counties, reflecting a diverse assemblage of angiosperm taxa detected across samples.

### Variation in plant volatile profiles

To characterize variation in odor chemistry among candidate mosquito-associated plants, we collected headspace samples from 42 plant taxa and analyzed them using GC-MS. This set included taxa that overlapped with mosquito-associated plant DNA surveys where possible, as well as plants selected because of prior evidence of mosquito attraction, abundance at field sites, or floral and vegetative traits consistent with mosquito-visited resources (Peach and Gries, 2019, 2020; Shannon et al., 2024; Upshur et al., 2023). Across 168 plant headspace collections, GC-MS analyses detected 211 volatile compounds. Volatile richness ranged from 12 to 96 compounds per taxon, and compound richness differed significantly among plant groups (Kruskal-Wallis H = 122.04, p < 0.001). Mean total volatile emission rates also varied strongly among plant groups, ranging from 0.312 to 159.71 ng h⁻¹. Emission rates differed significantly among plant groups (Kruskal-Wallis H = 109.53, p < 0.001), and plant-group compound richness was positively associated with mean total emission rate (Spearman ρ = 0.54, p < 0.001). Mean emissions were highest in groups including *Erigeron annuus*, *Solidago canadensis*, *Coreopsis tinctoria*, *Eupatorium capillifolium*, *Phlox* sp., and *Bidens alba*, whereas lower-emitting groups included *Sorghum* sp., *Tecoma stans*, *Senna alata*, *Paspalum notatum*, and *Lepidium virginicum* (Table S1).

Despite this quantitative variation, several compounds were broadly distributed and, in some plant groups, made up substantial proportions of the odor blend. For instance, 4-ethylacetophenone, α-pinene, and limonene were each detected in at least half of the replicates for 41 plant groups, reaching mean proportional abundances of 54.3% in *Plumbago auriculata*, 25.9% in *Tecoma stans*, and 23.1% in *Solidago canadensis*, respectively. Other broadly distributed compounds included 2-ethyl-1-hexanol, which was detected in at least half of the replicates for 27 plant groups and reached 28.9% in *Salvia splendens*, and 4-ethylbenzaldehyde, which was detected in at least half of the replicates for 29 plant groups and reached 19.3% in *Sorghum* sp. Caryophyllene was less ubiquitous but still common, occurring in at least half of the replicates for 22 plant groups and exceeding 5% mean abundance in 11 plant groups. Several of these broadly distributed compounds belong to volatile classes previously implicated in mosquito responses to plant-derived odors, including terpenes, aldehydes, alcohols, and ketones (Nyasembe and Torto, 2014; Sobhy and Berry, 2024).

However, broad compound sharing did not necessarily translate into similar odor blends, as illustrated by taxa with contrasting emission profiles. Among *Asclepias curassavica*, *Bidens alba*, and *Lantana camara*, mean total emission was highest in *B. alba* (36.5 ng h⁻¹), followed by *A. curassavica* (25.3 ng h⁻¹) and *L. camara* (15.2 ng h⁻¹), whereas volatile richness was greatest in *L. camara* and *B. alba* (96 and 90 detected compounds, respectively) compared with *A. curassavica* (51 compounds). Their dominant chemistry also differed sharply: *A. curassavica* was characterized by methyl salicylate and ethyl salicylate, *B. alba* by monoterpenes including α-pinene, β-myrcene, and β-pinene, and *L. camara* by sesquiterpenes including caryophyllene, α-zingiberene, germacrene D, and (E)-β-farnesene. Although 28 compounds were shared by all three taxa, these shared compounds often occurred at very different proportions, including methyl salicylate, α-pinene, and caryophyllene. Thus, overlap in compound presence did not necessarily correspond to similar odor blend structure.

We next focused on the five field-associated taxa selected for semi-field behavioral assays: *Bidens alba*, *Lantana camara*, *Richardia scabra*, *Solanum lycopersicum*, and *Sorghum* sp. These taxa spanned five plant families and provided a focused comparison of candidate mosquito-associated sugar hosts relevant to mosquito foraging. Total volatile emission rates differed significantly among these five taxa (Kruskal-Wallis H = 30.82, p < 0.001), ranging from low-emitting *Sorghum* sp. to higher-emitting *B. alba* and *L. camara*. Their volatile richness also differed, with 20 detected compounds in *Sorghum* sp., 45 in *S. lycopersicum*, 63 in *R. scabra*, 90 in *B. alba*, and 96 in *L. camara*. Even so, 17 compounds were detected in all five taxa and 28 compounds were detected in at least four taxa. The five profiles also differed in dominant compounds: *B. alba* was enriched in α-pinene, β-myrcene, and β-pinene; *L. camara* in caryophyllene, α-zingiberene, cis-β-ocimene, germacrene D, and (E)-β-farnesene; *R. scabra* in 4-ethylacetophenone, 2-ethyl-1-hexanol, α-pinene, limonene, and ethyl benzoate; *S. lycopersicum* in β-phellandrene; and *Sorghum* sp. in 4-ethylacetophenone, 2-ethyl-1-hexanol, 4-ethylbenzaldehyde, limonene, and α-pinene (Figure 1A).

**Figure 1.**
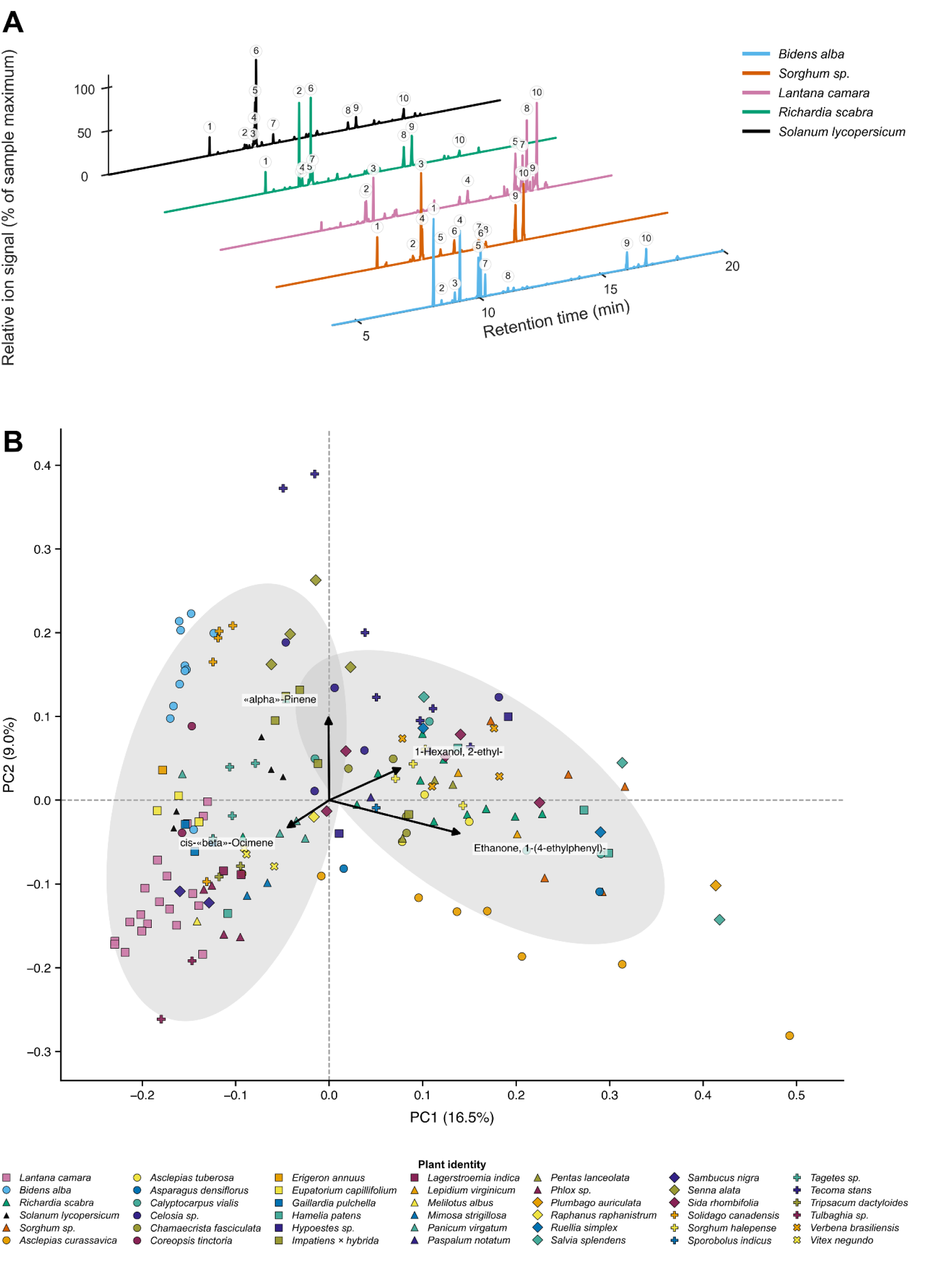
**(A).** Chromatograms of representative headspace volatile profiles from the five plant species used in the semi-field behavioral assay. Each trace is normalized to its own maximum. Numbered labels correspond to the top 10 compounds for each trace. **Label key: *Bidens alba***: 1, α-Pinene; 2, Camphene; 3, β-Pinene; 4, β-Myrcene; 5, Limonene; 6, trans-β-Ocimene; 7, cis-β-Ocimene; 8, Linalool; 9, Caryophyllene; 10, Germacrene D. ***Sorghum* sp.**: 1, α-Pinene; 2, 3-Carene; 3, 2-ethyl-1-hexanol; 4, Limonene; 5, Dihydromyrcenol; 6, Nonanal; 7 and 8, 4-ethylbenzaldehyde; 9 and 10, 4-ethylacetophenone. ***Lantana camara***: 1, 2-ethyl-1-hexanol; 2, Limonene; 3, cis-β-Ocimene; 4, 4-ethylacetophenone; 5, Caryophyllene; 6, cis-Geranylacetone; 7, cis-β-Farnesene; 8, Humulene; 9, 1-(1,5-dimethylhex-4-en-1-yl)-4-methylbenzene; 10, α-Zingiberene. ***Richardia scabra***: 1, α-Pinene; 2, (Z)-3-Hexenyl acetate; 3, 3-Carene; 4, hexyl acetate; 5, 2-ethyl-1-hexanol; 6, Limonene; 7, Eucalyptol; 8 and 9, 4-ethylacetophenone; 10, Caryophyllene. ***Solanum lycopersicum***: 1, α-Pinene; 2, α-Phellandrene; 3, o-Cymene; 4, 2-ethyl-1-hexanol; 5, Limonene; 6, β-Phellandrene; 7, Dihydromyrcenol; 8 and 9, 4-ethylacetophenone; 10, Caryophyllene. ***(B).*** Principal component analysis (PCA) of relative volatile compound proportions from plant headspace samples. Points represent individual samples, with colors and symbols indicating plant identity. Grey ellipses show the two descriptive k-means clusters, and vectors indicate the four compounds contributing most strongly to variation along the PCA axes.

The compositional differences between all 42 sampled taxa were reflected in multivariate analyses of relative volatile abundances (Figure 1B). Although several plant groups occupied overlapping regions of ordination space, volatile composition differed significantly among taxa (PERMANOVA: pseudo-F = 6.713, p = 0.001; ANOSIM: R = 0.751, p = 0.001). These differences were not attributable to differences in multivariate dispersion (PERMDISP: F = 2.544, p = 0.995).

### Semi-field behavioral assays

To evaluate whether field-associated plants elicited mosquito attraction under semi-field conditions, we used sticky traps arranged inside screen house assay arenas to compare plant-baited treatments with controls (Figure 2A). Assays focused on five taxa chosen from preliminary surveys of mosquito-associated plant DNA and common field-site vegetation: *Bidens alba, Lantana camara, Richardia scabra, Solanum lycopersicum*, and *Sorghum sp*.

**Figure 2.**
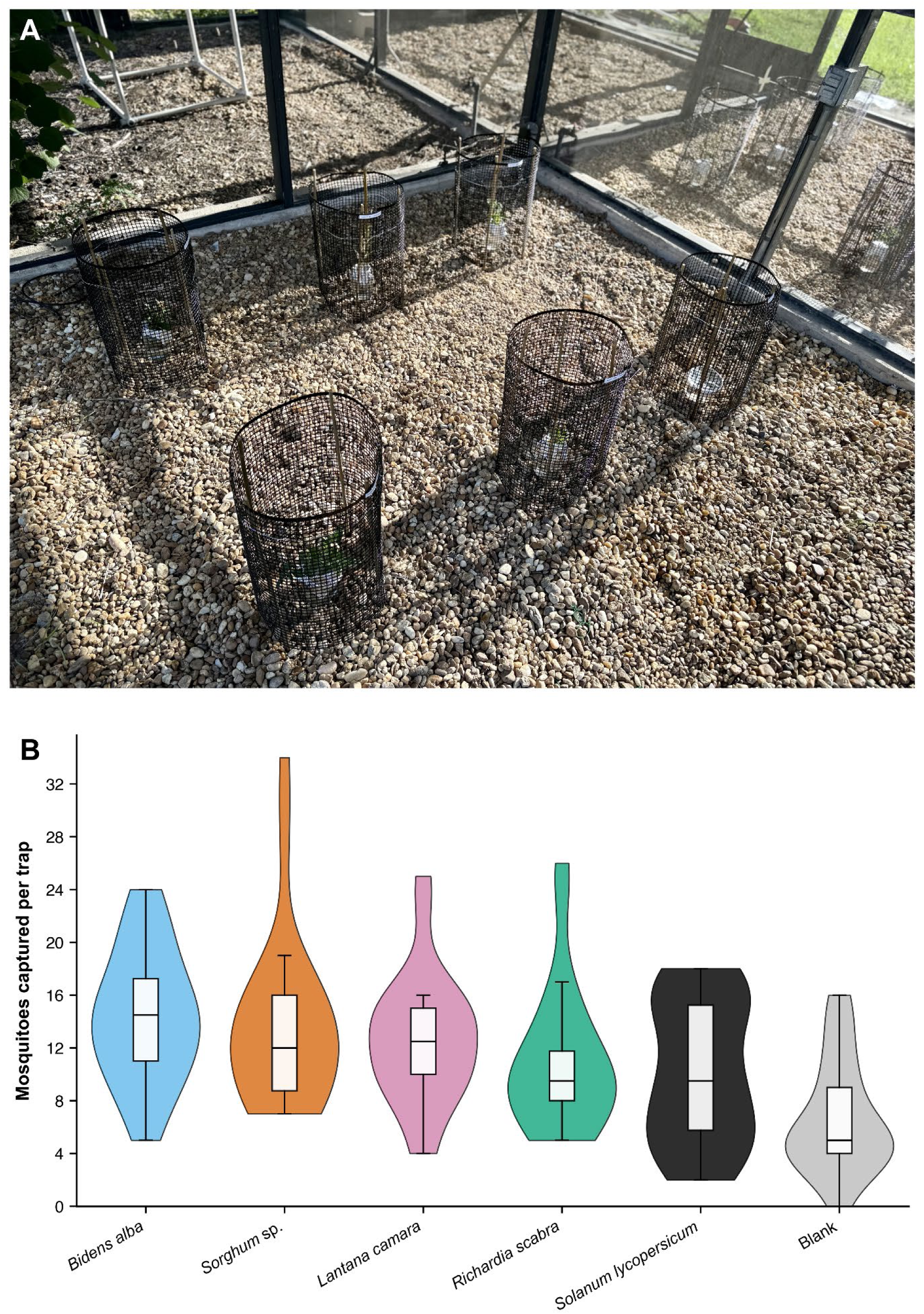
**(A).** Sticky trap arrangement used in the semi-field behavioral assay. Plant treatments and blank controls were placed within cylindrical mesh enclosures inside screen house assay arenas, with sticky traps used to quantify mosquito captures during each trial. ***(B).*** Distribution of mosquito captures per trap across plant treatments in semi-field sticky trap assays. Violin plots show the number of mosquitoes captured per trap for each plant species and the blank control across all trials. Embedded boxplots indicate medians and interquartile ranges.

Together, these taxa spanned five plant families and provided a focused set of candidate mosquito-associated plant taxa. Mosquito capture differed across plant treatments, with plant-baited traps capturing more mosquitoes on average than controls (Figure 2B). Mean trap catches were highest for *B. alba* (14.4 ± 1.0 mosquitoes per trap, mean ± SEM), followed by *Sorghum sp.* (13.6 ± 1.3) and *L. camara* (12.6 ± 1.0). *R. scabra* and *S. lycopersicum* showed intermediate mean catches of 11.0 ± 1.1 and 10.2 ± 1.1 mosquitoes per trap, respectively, whereas the blank control had the lowest mean catch (6.8 ± 0.9). These differences were supported by a negative binomial model with plant treatment as a fixed effect and day included as a blocking term (likelihood-ratio χ²(5) = 53.56, p < 0.001). Relative to the blank control, all plant treatments yielded significantly higher captures after Bonferroni correction. The strongest effects were observed for *B. alba* (IRR = 2.19, 95% CI = 1.75-2.73, adjusted p < 0.001), *Sorghum sp.* (IRR = 2.03, 95% CI = 1.63-2.53, adjusted p < 0.001), and *L. camara* (IRR = 1.88, 95% CI = 1.50-2.35, adjusted p < 0.001), followed by *R. scabra* (IRR = 1.64, 95% CI = 1.30-2.06, adjusted p < 0.001) and *S. lycopersicum* (IRR = 1.54, 95% CI = 1.22-1.93, adjusted p = 0.00137). A secondary weather-adjusted GLMM indicated that ambient conditions contributed to day-to-day variation in trap capture, but did not alter the main treatment pattern: including day-level weather improved model fit (χ²(3) = 18.55, p < 0.001), while plant treatment remained strongly significant after weather adjustment (χ²(5) = 50.90, p < 0.001).

To assess whether mosquitoes engaged in sugar feeding within the assay environment, mosquitoes collected from BG traps at the end of selected trials were analyzed using a warm anthrone assay. Across sampled days, roughly one-quarter of mosquitoes were classified as sugar-fed, indicating that mosquitoes fed on sugar sources during the semi-field assays. Because anthrone assays were performed on mosquitoes collected from BG traps rather than sticky traps, these data confirmed sugar feeding within the assay system but did not identify which plant taxa provided the sugar meal.

Finally, because the five tested taxa differed in volatile output, we explored whether species-level mean total volatile emission rates from GC-MS headspace collections were associated with mosquito capture in the semi-field assays. This species-level analysis was not significant (Pearson r = 0.61, exact permutation p = 0.292; Spearman ρ = 0.30, exact permutation p = 0.683). Notably, *Sorghum sp*. had the lowest mean volatile emission rate among the five taxa but the second-highest mean trap capture, indicating that total volatile output alone did not explain the behavioral pattern.

## Discussion

While molecular approaches have begun to reveal the diversity of plant taxa associated with mosquitoes in nature, considerably less is known about whether these field-associated taxa correspond to behaviorally relevant resources, or what chemical features characterize them. To address these questions, we combined plant DNA barcoding, semi-field behavioral assays, and volatile profiling to examine mosquito–plant interactions across multiple levels of biological organization. Our results suggest that *Aedes aegypti* utilizes a broad diversity of plant taxa in nature and that some of these field-associated taxa can also elicit attraction under semi-field conditions. DNA barcoding revealed associations with 90 genera across 37 plant families in three Florida counties, and representative taxa selected from these field associations were shown to consistently attract more mosquitoes than blank controls in semi-field behavioral assays. Furthermore, volatile profiling across 42 plant taxa revealed a chemically diverse odor landscape characterized by both broadly shared compounds and substantial variation in odor blend composition among species. Similar patterns were evident among the five taxa evaluated in behavioral assays, which exhibited considerable overlap in volatile composition despite belonging to different plant families.

Recent molecular studies have uncovered mosquito-plant associations spanning diverse plant taxa. For example, Nyasembe et al. (2018) used DNA barcoding to identify plants associated with field-collected *Aedes* and *Anopheles* mosquitoes in Kenya, then analyzed volatiles from the identified plants to ask whether host plants shared common chemical features. They identified mosquito species-specific associations with taxa including *Pithecellobium*, *Senna*, *Opuntia*, *Ricinus*, and *Leonotis*. Despite these differences in host-plant use, several volatile compounds, including β-myrcene and (E)-β-ocimene, were detected across multiple host plants, suggesting that both host-plant specificity and shared plant-odor features may shape mosquito plant seeking (Nyasembe et al., 2018). Kinya et al. (2024) similarly used DNA barcoding to identify plant material associated with wild Kenyan anophelines and found that detections were dominated by acacia and other Fabaceae taxa. However, because those detections were not paired with plant availability data, the authors cautioned that the pattern could not necessarily be interpreted as plant preference (Kinya et al., 2024). In another recent study in Florida, Mosore et al. (2026) used rbcL metabarcoding to identify plants associated with *Culex quinquefasciatus* across six counties and identified a broad range of taxa, including cultivated plants, non-cultivated plants, and grasses. Their results broaden the known plant associations of *Cx. quinquefasciatus*, but also highlight an important interpretive challenge: that plant DNA detections may reflect nectar feeding, but may also reflect the use of other plant-derived sugar sources such as honeydew or guttation droplets, or environmental contamination (Mosore et al., 2026). Consequently, these studies demonstrate that molecular approaches are indeed powerful for generating ecologically grounded candidate plant associations, but that plant DNA detections alone do not establish attraction, preference, or resource value. Molecular identifications are perhaps most informative when combined with complementary behavioral, chemical, or ecological data that can help place detected taxa into a broader biological context.

Beyond documenting broad taxonomic diversity, molecular surveys can also reveal recurring associations that may help identify particularly relevant taxa for downstream investigation. In our data, plant detections were not uniformly distributed across taxa. Instead, some genera were detected repeatedly or were especially prominent within particular counties, suggesting that *Ae. aegypti* may utilize a wide range of plant resources while also repeatedly interacting with a subset of locally available taxa. Similar unevenness has emerged in other mosquito–plant surveys, including the acacia-dominated detections reported in Kenyan anophelines and the recurring plant associations observed in Florida *Cx. quinquefasciatus* (Kinya et al., 2024; Mosore et al., 2026). In our study, *Datura* was frequently detected in St. Johns and Volusia counties, while genera such as *Bidens*, *Richardia*, and *Solanum* were also repeatedly detected and provided ecologically grounded candidates for behavioral and chemical follow-up. These recurring detections likely reflect a combination of ecological and methodological factors, including local plant availability, phenology, accessibility, mosquito behavior, and molecular detectability, but they nevertheless help distinguish repeated field associations from the broader background of plant diversity. Importantly, repeated detections do not by themselves establish attraction, but they provide a field-derived basis for selecting plant taxa for behavioral assays, an important next step given that mosquitoes can differ in their attraction to plant-derived resources (Müller et al., 2011; Sissoko et al., 2019). In this sense, repeated detections are valuable as a bridge between molecular surveys and experimental assays, identifying plant taxa that can subsequently be evaluated for attraction, behavioral preference, and odor chemistry.

Mosquitoes are known to respond to a wide variety of plant-associated sensory cues, including floral odors, fruit volatiles, vegetative emissions, and other sugar-source-associated cues such as honeydew (Foster and Hancock, 1994; Foster, 1995; Pullmann-Lindsley et al., 2024). Numerous laboratory, semi-field, and field studies have shown that mosquitoes do not respond equally to all plant-derived resources, with attraction varying among plant species, mosquito species, sex, physiological state, and assay context (Müller et al., 2011; Nikbakhtzadeh et al., 2014; Fikrig et al., 2017; Sissoko et al., 2019; Meza et al., 2020; Kashiwagi et al., 2022). For instance, Sissoko et al. (2019) found that *Ae. aegypti* fed readily on several plant-derived sugar sources in laboratory assays, but field traps revealed differential attraction of males and females to only subsets of those resources, indicating that feeding opportunity and attraction are related but separable. Similarly, Müller et al. (2011) tested *Ae. albopictus* responses to flowers, fruits, seedpods, and honeydew-soiled foliage in the field and found that several flowers and fruits were attractive, whereas honeydew-soiled plants were not significantly attractive. Upshur et al. (2023) similarly showed that both *Ae. aegypti* and *Ae. albopictus* landed on and sugar-fed from several ornamental plant species, and paired those feeding assays with headspace GC-MS to characterize the plant odor profiles associated with those sugar resources. These studies demonstrate that plant-derived resources can influence mosquito behavior, but they provide limited insight into whether the plants tested are the same taxa that mosquitoes regularly encounter under natural conditions.

One of the strengths of molecular approaches is that they provide a means of identifying plant taxa associated with mosquitoes in nature. However, relatively few studies have carried these field-derived associations forward into downstream behavioral or chemical assays. In the present study, we selected five plant taxa for semi-field behavioral assays because they were identified in preliminary mosquito barcoding surveys, were common at field sites, and represented a taxonomically diverse subset of candidate sugar hosts spanning five plant families. All five plant treatments attracted more mosquitoes than blank controls, providing supporting evidence that taxa associated with mosquitoes in nature can also function as behaviorally relevant cues under semi-field conditions. Anthrone assays confirmed that mosquitoes were sugar feeding within the semi-field environment, supporting the biological relevance of the assay, although they do not identify which plant provided the sugar meal. Thus, these taxa were not intended to represent the most important mosquito-associated plants overall, but rather a field-grounded set of candidates for evaluating how naturally encountered plants influence mosquito orientation. At the same time, the observation that all plant treatments outperformed blank controls suggests that *Ae. aegypti* can respond to multiple field-associated plant taxa under semi-field conditions. Notably, total volatile emission rate was not significantly associated with mosquito captures among the five tested taxa, suggesting that attraction was not simply a function of producing more odor. This pattern fits with broader floral-scent and mosquito chemical-ecology work showing that behavioral responses often depend on blend composition, compound ratios, and sensory context rather than total odor abundance alone (Raguso, 2008; Nyasembe et al., 2012; Lahondère et al., 2020; Meza et al., 2020). For instance, Nyasembe et al. (2012) showed that female *Anopheles gambiae* attraction to plant-derived volatile mixtures depended on the relative concentrations of antennally active compounds, with optimized synthetic mixtures outperforming both intact plant odor and blends mimicking natural proportions. Meza et al. (2020) similarly found that responses to mango-bait volatiles depended on compound identity and blend context: some individual compounds elicited attraction, myrcene elicited avoidance, and a three-component blend was strongly attractive. In an orchid-associated system, Lahondère et al. (2020) showed that mosquito attraction depended on odor-blend composition, particularly the balance between nonanal-rich attractive scent and lilac aldehyde-rich scents from related orchids. Thus, mosquito responses to the five plants tested here may depend on blend structure, relative abundances of particular compounds, whole-plant context, or interactions among olfactory, visual, and other sensory cues.

Plant odors are highly diverse, often comprising complex odors of dozens to hundreds of volatile compounds that vary substantially among species, tissues, phenological stages, and ecological contexts (Raguso, 2008; Sobhy and Berry, 2024). At the same time, many plant-derived volatiles recur across taxonomically diverse plants, raising the possibility that mosquitoes encounter both taxon-specific odor signatures and a shared chemical background while foraging. Reviews of mosquito–plant semiochemistry have emphasized that recurring plant-derived compounds occur across diverse taxa, although their biological activity depends on concentration, blend context, and the mosquito species receiving the signal (Nyasembe and Torto, 2014; Sobhy and Berry, 2024). Together, these studies suggest that plant odors may be characterized by both shared and taxon-specific chemical features. Our results were broadly consistent with this view. Volatile profiling across 42 plant taxa revealed a chemically diverse odor landscape, yet many compounds were broadly distributed throughout the dataset. Compounds such as α-pinene, limonene, 4-ethylacetophenone, 2-ethyl-1-hexanol, 4-ethylbenzaldehyde, and caryophyllene occurred across numerous taxa and, in some cases, represented substantial portions of individual odor profiles. At the same time, multivariate analyses indicated significant differences in volatile composition among plant taxa, reflecting substantial variation in odor composition despite the presence of shared compounds. Thus, the plant taxa associated with mosquitoes in this study should be viewed as chemically distinct odor sources embedded within a broader plant-associated volatile background.

The five plant taxa evaluated in semi-field behavioral assays illustrate this pattern particularly well. Although these species represented five plant families and differed in volatile composition, 17 compounds were detected in all five taxa and 28 compounds were shared among at least four species. Similar combinations of overlap and divergence have been observed in other mosquito–plant systems. Jhumur et al. (2008) found that *Culex pipiens molestus* was attracted to floral odors from geographically distinct populations of *Silene otites*, even though attraction was not explained by total scent emission and electrophysiological responses did not simply track the most abundant compounds in the floral odors. Yu et al. (2015) similarly showed that female *Culex pipiens pallens* responded to common host-plant volatiles in a concentration-dependent manner, with many compounds attractive only within particular dose ranges and a reduced synthetic blend performing as well as a larger blend. In an orchid-associated system, Lahondère et al. (2020) further showed that mosquito attraction depended on the relative balance of floral odor components rather than the presence of individual compounds alone. Collectively, these findings suggest that mosquito attraction may depend less on individual compounds in isolation than on the broader composition, concentration, and structure of odors. This interpretation is also consistent with our behavioral results. Although all five plant taxa increased mosquito captures relative to blank controls, total volatile emission rate was not significantly associated with mosquito attraction. *Sorghum* sp., for example, produced comparatively low total volatile emissions yet remained attractive under semi-field conditions. These observations suggest that attraction is not simply a function of odor quantity, but may instead depend on blend composition, relative abundances of particular compounds, or interactions among olfactory, visual, and other sensory cues.

Whether these patterns are characteristic of mosquito-associated plants more generally remains an open question. Although plant associations were characterized across three Florida counties, broader geographic sampling will be needed to determine whether similar patterns emerge across regions, seasons, and plant communities. Likewise, while county-level plant surveys provided an important framework for interpreting molecular detections, finer-scale measurements of plant availability, flowering phenology, and local habitat composition could help clarify the extent to which recurring detections reflect resource availability versus repeated mosquito interactions with particular taxa. Additional behavioral assays involving a larger set of recurrently detected plants would further help determine whether some field-associated taxa consistently elicit stronger attraction than others. More broadly, future studies that further integrate molecular surveys, behavioral assays, and volatile analyses across a wider range of plant taxa may help clarify which features of naturally occurring plant odor sources are most relevant to mosquito plant seeking. Such efforts could strengthen links between field-derived plant associations and the sensory cues that guide mosquito foraging, while providing a more complete picture of mosquito–plant interactions in natural environments.

## Acknowledgments

We are grateful for the discussions and advice provided by A. Rouyar, C. Ruiz, O. Akbari, and J. Pitts. We thank B. Nguyen for mosquito rearing and care.

## Author contributions

Conceptualization: SJ and JAR; Methodology: SJ, AP, MTM, ERB, and JAR; Formal analysis: SJ and JAR; Investigation: SJ, AP, and JAR; Data curation: SJ and JAR; Writing - review & editing: SJ, AP, MSM, ERB, and JAR; Visualization: SJ; Supervision: JAR; Project administration: JAR; Funding acquisition: JAR.

## Data availability

Code is available at https://github.com/riffelllab.

## Supplementary data

***Table S1.*** Compound emission rates estimated from GC-MS headspace collections across plant groups. The first row reports mean total emission rate for each plant group, standardized to the 12-h collection period and reported as ng h⁻¹; subsequent rows report each compound’s proportional contribution to that total. Dashes indicate compounds not detected in a given plant group.

## References

Barredo, E., & DeGennaro, M. (2020). Not Just from Blood: Mosquito Nutrient Acquisition from Nectar Sources. Trends in Parasitology, 36(5), 473–484. 10.1016/j.pt.2020.02.003

Cooper, A. N., Malmgren, L., Hawkes, F. M., Farrell, I. W., Hien, D. F. d S., Hopkins, R. J., Lefèvre, T., & Stevenson, P. C. (2025). Identifying mosquito plant hosts from ingested nectar secondary metabolites. Scientific Reports, 15(1), 6488. 10.1038/s41598-025-88933-1

Fikrig, K., Johnson, B. J., Fish, D., & Ritchie, S. A. (2017). Assessment of synthetic floral-based attractants and sugar baits to capture male and female Aedes aegypti (Diptera: Culicidae). Parasites & Vectors, 10(1), 32. 10.1186/s13071-016-1946-y

Fikrig, K., Peck, S., Deckerman, P., Dang, S., Fleur, K. S., Goldsmith, H., Qu, S., Rosenthal, H., & Harrington, L. C. (2020). Sugar feeding patterns of New York Aedes albopictus mosquitoes are affected by saturation deficit, flowers, and host seeking. PLOS Neglected Tropical Diseases, 14(10), e0008244. 10.1371/journal.pntd.0008244

Foster, W. A. (1995). Mosquito sugar feeding and reproductive energetics. Annual Review of Entomology, 40, 443–474. 10.1146/annurev.en.40.010195.002303

Foster, W. A., & Hancock, R. G. (1994). Nectar-related olfactory and visual attractants for mosquitoes. Journal of the American Mosquito Control Association, 10(2 Pt 2), 288–296.

Jhumur, U. S., Dötterl, S., & Jürgens, A. (2008). Floral Odors of Silene otites: Their Variability and Attractiveness to Mosquitoes. Journal of Chemical Ecology, 34(1), 14–25. 10.1007/s10886-007-9392-0

Jové, V., Gong, Z., Hol, F. J. H., Zhao, Z., Sorrells, T. R., Carroll, T. S., Prakash, M., McBride, C. S., & Vosshall, L. B. (2020). Sensory Discrimination of Blood and Floral Nectar by Aedes aegypti Mosquitoes. Neuron, 108(6), 1163–1180.e12. 10.1016/j.neuron.2020.09.019

Kashiwagi, G. A., Oppen, S. von, Harburguer, L., & González-Audino, P. (2022). The main component of the scent of Senecio madagascariensis flowers is an attractant for Aedes aegypti (L.) (Diptera: Culicidae) mosquitoes. Bulletin of Entomological Research, 112(6), 837–846. 10.1017/S0007485322000256

Kinya, F., Milugo, T. K., Mutero, C. M., Wondji, C. S., Torto, B., & Tchouassi, D. P. (2024). Insights into malaria vectors–plant interaction in a dryland ecosystem. Scientific Reports, 14(1), 20625. 10.1038/s41598-024-71205-9

Lahondère, C., Vinauger, C., Okubo, R. P., Wolff, G. H., Chan, J. K., Akbari, O. S., & Riffell, J. A. (2020). The olfactory basis of orchid pollination by mosquitoes. Proceedings of the National Academy of Sciences, 117(1), 708–716. 10.1073/pnas.1910589117

Manda, H., Gouagna, L. C., Foster, W. A., Jackson, R. R., Beier, J. C., Githure, J. I., & Hassanali, A. (2007). Effect of discriminative plant-sugar feeding on the survival and fecundity of Anopheles gambiae. Malaria Journal, 6(1), 113. 10.1186/1475-2875-6-113

Meza, F. C., Roberts, J. M., Sobhy, I. S., Okumu, F. O., Tripet, F., & Bruce, T. J. A. (2020). Behavioural and Electrophysiological Responses of Female Anopheles gambiae Mosquitoes to Volatiles from a Mango Bait. Journal of Chemical Ecology, 46(4), 387–396. 10.1007/s10886-020-01172-8

Mosore, M.-T., Mishra, S., Villa, M., Agbodzi, B., Estep, A. S., Prasauskas, A., Qualls, W. A., Killingsworth, D., Unlu, I., Tressler, M., Dinglasan, R. R., & Burgess, E. R. (2026). A Tandem Metabarcoding and Taxonomic Forensics Approach to Surveillance of Mosquito–Plant Interactions for Culex quinquefasciatus in Florida. Insects, 17(1), 13. 10.3390/insects17010013

Müller, G. C., Xue, R.-D., & Schlein, Y. (2011). Differential attraction of Aedes albopictus in the field to flowers, fruits and honeydew. Acta Tropica, 118(1), 45–49. 10.1016/j.actatropica.2011.01.009

Nikbakhtzadeh, M. R., Terbot, J. W., Otienoburu, P. E., & Foster, W. A. (2014). Olfactory basis of floral preference of the malaria vector Anopheles gambiae (Diptera: Culicidae) among common African plants. Journal of Vector Ecology, 39(2), 372–383. 10.1111/jvec.12113

Njoroge, T. M., Calla, B., Berenbaum, M. R., & Stone, C. M. (2021). Specific phytochemicals in floral nectar up-regulate genes involved in longevity regulation and xenobiotic metabolism, extending mosquito life span. Ecology and Evolution, 11(12), 8363–8380. 10.1002/ece3.7665

Nyasembe, V. O., Tchouassi, D. P., Pirk, C. W. W., Sole, C. L., & Torto, B. (2018). Host plant forensics and olfactory-based detection in Afro-tropical mosquito disease vectors. PLOS Neglected Tropical Diseases, 12(2), e0006185. 10.1371/journal.pntd.0006185

Nyasembe, V. O., Teal, P. E., Mukabana, W. R., Tumlinson, J. H., & Torto, B. (2012). Behavioural response of the malaria vector Anopheles gambiae to host plant volatiles and synthetic blends. Parasites & Vectors, 5(1), 234. 10.1186/1756-3305-5-234

Nyasembe, V. O., & Torto, B. (2014). Volatile phytochemicals as mosquito semiochemicals. Phytochemistry Letters, 8, 196–201. 10.1016/j.phytol.2013.10.003

Peach, D. A. H., & Gries, G. (2020). Mosquito phytophagy—Sources exploited, ecological function, and evolutionary transition to haematophagy. Entomologia Experimentalis et Applicata, 168(2), 120–136. 10.1111/eea.12852

Peach, D., & Gries, G. (2019). Supplementary data for: Mosquito phytophagy—Sources exploited, ecological function, and evolutionary transition to haematophagy (Version 4) [Dataset]. Dryad. 10.5061/DRYAD.63XSJ3TZ5

Pullmann-Lindsley, H., Huff, R. M., Boyi, J., & Pitts, R. J. (2024). Odorant receptors for floral-and plant-derived volatiles in the yellow fever mosquito, Aedes aegypti (Diptera: Culicidae). PLOS ONE, 19(5), e0302496. 10.1371/journal.pone.0302496

Raguso, R. A. (2008). Wake Up and Smell the Roses: The Ecology and Evolution of Floral Scent. Annual Review of Ecology, Evolution, and Systematics, 39, 549–569. 10.1146/annurev.ecolsys.38.091206.095601

Shannon, D. M., Richardson, N., Lahondère, C., & Peach, D. (2024). Mosquito floral visitation and pollination. Current Opinion in Insect Science, 65, 101230. 10.1016/j.cois.2024.101230

Sissoko, F., Junnila, A., Traore, M. M., Traore, S. F., Doumbia, S., Dembele, S. M., Schlein, Y., Traore, A. S., Gergely, P., Xue, R.-D., Arheart, K. L., Revay, E. E., Kravchenko, V. D., Beier, J. C., & Müller, G. C. (2019). Frequent sugar feeding behavior by Aedes aegypti in Bamako, Mali makes them ideal candidates for control with attractive toxic sugar baits (ATSB). PLOS ONE, 14(6), e0214170. 10.1371/journal.pone.0214170

Sobhy, I. S., & Berry, C. (2024). Chemical ecology of nectar–mosquito interactions: Recent advances and future directions. Current Opinion in Insect Science, 63, 101199. 10.1016/j.cois.2024.101199

Stone, C. M., Jackson, B. T., & Foster, W. A. (2012). Effects of Plant-Community Composition on the Vectorial Capacity and Fitness of the Malaria Mosquito Anopheles gambiae. The American Journal of Tropical Medicine and Hygiene, 87(4), 727–736. 10.4269/ajtmh.2012.12-0123

Upshur, I. F., Fehlman, M., Parikh, V., Vinauger, C., & Lahondère, C. (2023). Sugar feeding by invasive mosquito species on ornamental and wild plants. Scientific Reports, 13(1), 22121.

Yu, B.-T., Ding, Y.-M., & Mo, J.-C. (2015). Behavioural response of female Culex pipiens pallens to common host plant volatiles and synthetic blends. Parasites & Vectors, 8(1), 598. 10.1186/s13071-015-1212-8

